# Fluctuation, correlation and perturbation-response behavior of nature-made and artificial nanobodies

**DOI:** 10.1101/2020.02.06.936856

**Authors:** Aysima Hacisuleyman, Batu Erman, Albert Erkip, Burak Erman

**Affiliations:** Department of Chemical and Biological Engineering, Koc University, Sariyer 34450, Istanbul, Turkey; Molecular Biology, Genetics and Bioengineering Program, Faculty of Engineering and Natural Sciences, Sabanci University, Tuzla 34956, Istanbul, Turkey; Faculty of Engineering and Natural Sciences, Sabanci University, Tuzla 34956, Istanbul, Turkey

## Abstract

Nanobodies, like other antibodies bind their targets through complementarity determining regions (CDR’s). Improving nanobody-antigen binding affinities by introducing mutations in these CDR’s is critical for biotechnological applications. However, any mutation is expected to introduce changes in the behavior of the protein, such as fluctuations of residues, correlation of fluctuations of residue pairs, response of a residue to perturbation of another. Most importantly, the nanoscale dynamics of the protein may change. In these respects, the problem is similar to the general problem of dynamic allostery, a perturbation at one site affecting the response at another site. Using the Gaussian Network Model of proteins, we show that CDR mutations indeed modify the fluctuation profile and dynamics of the nanobody. Effects are not confined to CDR regions but extend throughout the full structure. We introduce a computational scheme where fluctuations of a residue are perturbed by a force and response amplitude and response time of the remaining residues are determined. Response to a perturbation of a residue shows a synchronous and an asynchronous component. The model is used to quantify the effects of mutation on protein dynamics: highly perturbable residues and highly responsive residues of the nanobody are determined. Residues whose perturbation has no effect on protein behavior may also be determined with the present model. Three known nanobodies produced by nature are used as an illustrative example and their various modifications which we generated by CDR residue mutations are analyzed.

## Introduction

Nanobodies are single chain antibodies produced by members of Camelidae family (*Llama glama, Vicugna pacos, Camelus dromedaries, Camelidae, Camelus bactrianus*), nurse sharks, *Ginglymostoma cirratum*, wobbegongs, *Orectolobus* and spotted ratfish, *Hydrolagus colliei*. ^1^ They are highly specific and exert high affinity towards their target. They can be easily expressed in microorganisms for biochemical purification or intracellularly in target cells. Their toxicity is low, and their tissue penetration is not limited due their small size. The regions on nanobodies that are responsible for binding and recognition are called complementarity-determining regions (CDR).^1^ While the diversity of typical mammalian antibodies are derived from their heavy and light chain combinations, heavy chain antibodies (hcAbs) belong to the class of single domain antibodies (sdAbs). A single domain antibody is called V_H_H (variable heavy domain of heavy chain antibodies) domain also known as nanobody. Nanobodies contain complete antigen binding site. Due to their biochemical functionality and economic benefits, interest in nanobodies has grown in biotechnology and medicine. In nature, binding of hcAb proteins to their targets takes place through their three CDR loops. The millions of hcAb proteins that an animal can produce, differ in their amino acid sequence mainly in the CDR regions. This diversity is generated by a mutation inducing recombination process followed by selection of functional proteins in B lymphocytes. The natural mutational process can be mimicked in vitro by mutating the CDR of pre-existing nanobodies to alter or enhance affinity against alternative antigens, a solution that can be applied to drug design.^2^

There are, however, several problems in the generation of artificial nanobody drugs by modifying the natural ones by replacing residues on the CDRs. Nature selects the most appropriate nanobody for its target by successive mutations and successive elimination of the unsuitable ones. Each mutation causes changes in the structure and dynamics of the nanobody. If these changes improve the function of the nanobody, then it survives. Our knowledge of how a mutation will change the function of a nanobody is limited. Therefore, any new information pertinent to the nanobody function is valuable. The specific aim of this paper is to understand the effects of changing residues in the three CDR regions of a nanobody. Effects of replacing a residue with another may be divided into (i) inter and (ii) intramolecular factors. Intermolecular factors depend essentially on the physicochemical properties of the residue and the ones which will be in contact with it, including water molecules and spatially neighboring residues of the nanobody that are remote along its primary sequence. Potentials that describe intermolecular interactions are now well understood and well formulated.^3^ Intramolecular factors, as we show in the present paper, are important and deserve closer inspection. Among these, changes in the Ramachandran maps affecting the conformations of the residue and of first and second neighbors along the chain are the most dominant ones. Emphasis on the interactions of this type was introduced by Flory.^4^ Backbone conformations of a peptide unit are determined by two Ramachandran angles, phi and psi, and the bond angles of the peptide unit.^3, 4^ These angles are not fixed and exhibit fluctuations and lead to spatial fluctuations of the atoms of the chain appended to this peptide unit. Depending on the type of the peptide unit, some angles are confined to restricted regions while others may fluctuate more freely. Examples of tight and wide fluctuation regions are well documented in the work of Karplus ^5^ and the contributions of these fluctuations to configurational space of proteins were pointed out recently.^6^

Applications of nanobody technology require mutating CDR residues to make the protein more suitable (i.e. have higher affinity) for binding its target. These modifications could potentially lead to changes in conformations and dynamics of the remaining parts of the nanobody. Here, we show that upon mutation, fluctuations of residues not only in the CDR domain but in spatially remote parts of the nanobody, change significantly. These changes affect the correlations of residue fluctuations and the dynamic response of the protein to external perturbations.

Response of the protein to perturbation applied on certain residues is generally studied by linear response theory and focuses on relative changes in coordinates.^7-10^ In these studies, a constant force is applied on a residue and the resulting displacement of all other residues is calculated. In Ref. ^11^ the applied force is time dependent and is closest to the present study. The present work introduces changes in fluctuations and their time dependent correlations. The nanobody, like all other proteins, is a dynamic entity where each atom exhibits spatial fluctuations at characteristic times of nanoseconds. The extent of these fluctuations is measured by an average quantity, the mean square deviation (msd) of the atom. These are proportional to experimentally measurable Debye-Waller factors or the B-factors through the relation *B*_*i*_ = 8*π*^2^⟨(Δ*R*_*i*_)^2^⟩/ 3 where *B*_*i*_ is the experimentally measured B-factor, Δ*R*_*i*_ is the fluctuation of the i^th^ residue from its mean position and the term in angular brackets is the mean-square fluctuation of the i^th^ atom as will be discussed in detail below. The set of B-factors of all atoms, referred to as the ‘fluctuation profile’, is an intrinsic property, or a signature, of the protein resulting from the collective action of inter and intramolecular effects. Perturbation of a proteins by external forces and changes in its structure will change this signature.

Correlation of fluctuations of two atoms is another property built into the structure of the protein. This is defined as the projection of the instantaneous fluctuation of one atom on those of the other, averaged over time. Symbolically it is shown by the expression ⟨Δ*R*_*i*_(*t*)·Δ*R*_*j*_(*t*)⟩. The dot between the two vectors denotes the scalar product, and the angular brackets denote the time average. Two atoms are positively correlated if at each instant they move in the same direction or are negatively correlated if their fluctuations are in opposite directions. They are uncorrelated if the fluctuations of one are random relative to those of the other. Like the B-factors, correlations of atom pairs are also intrinsic properties of a protein and perturbation of the protein by external forces will change this signature. Two atoms can communicate with each other only if their fluctuations are correlated. Information, which is necessary for the protein to perform its function, cannot be transferred from one atom to another if their fluctuations are uncorrelated. In case the fluctuation of one atom is observed after a time delay of *τ*, then we obtain the time delayed correlation, *C*_*ij*_(*t,τ*) = ⟨Δ*R*_*i*_(*t*)·Δ*R*_*j*_(*t* + *τ*)⟩. This average quantity is obtained by looking at a large number of snapshots for atom i at time t, and of atom j at time *t* +*τ*, and averaging. This average shows us the effect of atom i on atom j and may be interpreted as the information transferred from atom i to j during the time interval*τ*. In a harmonic system, *C*_*ij*_(*t,τ*) = *C*_*ji*_(*t,τ*). Although correlation is a measure of information transfer, the more general definition of information transfer based on entropy may show that effect of i on j may not equal to that from j to i, even in harmonic systems.^12, 13^ As time *τ* increases, the effect of atom i on j averages out and goes to zero. *C*_*ij*_(*t,τ*) is a measure of information flow from i to j when the system fluctuates freely due to thermal noise. If atom i is perturbed, resulting from a mutation for example, then the correlation *C*_*ij*_(*t,τ*) changes. Existing correlations between a pair i and j may vanish and new correlations may emerge between a new pair, resulting in the change of function of the protein. The perturbation-response between atom pairs is also time dependent. The perturbation-response may be of long duration, or vanish rapidly. The duration of the effect of atom i on atom j is measured by the correlation time of the interaction. The correlation time of perturbation-response, which we introduce in this paper is of significant importance for understanding fast protein dynamics.^14^ One biotechnological application emerges when certain non-CDR residues of a nanobody are mutated to improve its penetration through the cell wall.^15^ For this reason, surface amino acids whose perturbation will not lead to a response have to be determined. Such nonresponsive residues may be determined with the proposed model.

Theoretical and computational models are convenient tools for understanding and predicting protein behavior. There are different levels of approximation in describing the behavior of the protein. The simplest one, given by the Gaussian Network Model (GNM),^16, 17^ is the harmonic approximation where all residues are attached to spatially neighboring residues by linear springs and fluctuate under the restoring forces coming from the others. We show that under external perturbation the residues of the protein fluctuate such that there is a synchronous as well as an asynchronous, out of phase response. Some residues respond to perturbations more than the others. This places emphasis on residues which induce the largest responses in the protein. Using the nanobody structure as an example and the harmonic approximation, we derive certain general rules on perturbation effects and show how they change with mutation.

We analyzed three different nanobodies produced in nature for binding to their specific targets. These nanobodies were crystallized in complex with their ligands, a) the extracellular domain of the human epidermal growth factor receptor (EGFR), b) human lysozyme and c) *Trypanosoma congolense* fructose-1,6-bisphosphate aldolase enzyme. The Protein Data Bank codes for the three nanobodies are 4KRO.pdb, 4I0C.pdb and 5O0W.pdb, respectively. We induced several mutations in the CDR regions of these proteins and compared the behavior of the mutated residues with the corresponding wild type nanobodies. In order to reduce the size of the paper, we present results for 4KRO here and give the results for 4I0C and 5O0W as Supplementary Material. In order to reduce cross referencing, the mathematics of the model is not delegated to the Supplementary part but presented in an Appendix at the end of the paper.

## Method

In a nanobody, the number of atoms is in the order of one thousand which is too high to address. For this reason, we adopt a coarse-grained approach and consider the correlations between residues. More specifically, we assume that the atoms of a residue are collapsed on its alpha carbon and study the correlations between alpha carbons of each residue. Throughout the paper, we use the term correlation of fluctuations of two residues synonymously with correlation of fluctuations of alpha carbons.

The nanobody structure EgA1 in complex with extracellular region of EGFR, obtained from Protein Data Bank with PDB id 4KRO and two modified nanobodies, named 4KROm1 and 4KROm2 obtained by mutating the CDR residues of EgA1 in 4KRO are studied. The ribbon diagram of 4KRO is shown in Figure 1.

**Figure 1.**
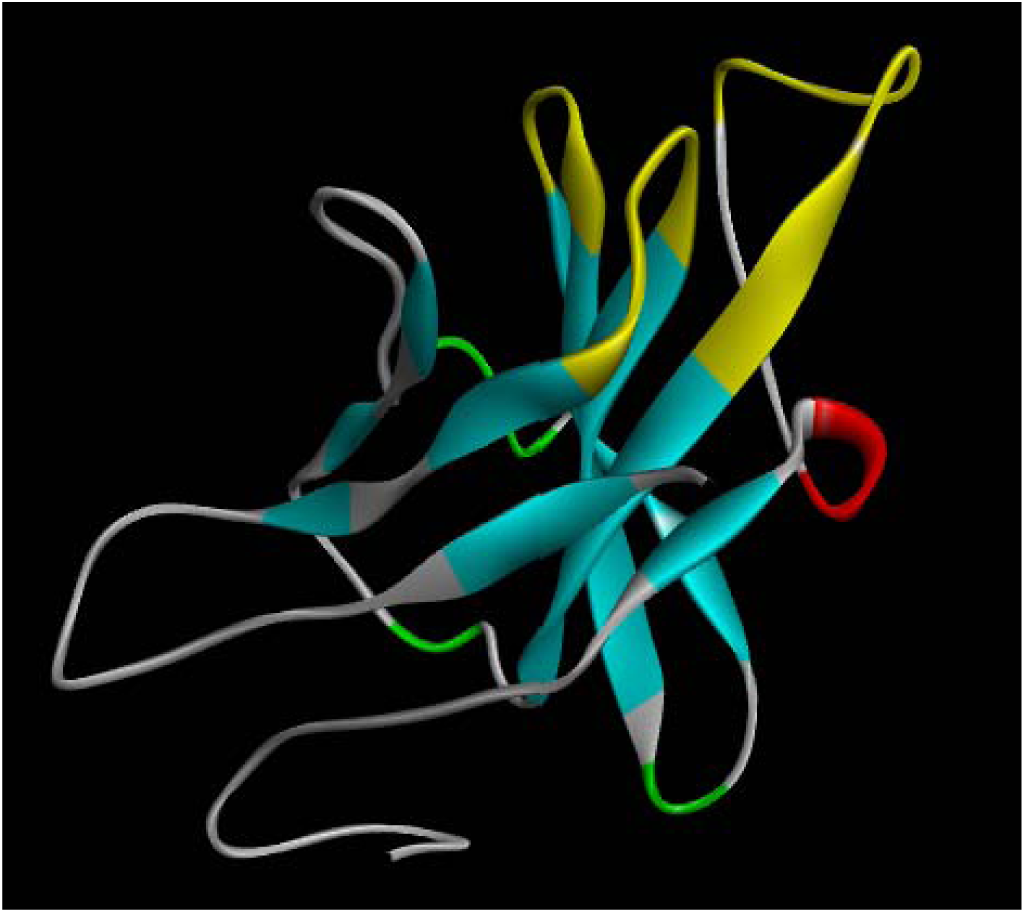
Ribbon diagram of the Nanobody/VHH domain EgA1 in 4KRO.pdb. The three CDR regions are highlighted in yellow. It has eight beta strands that stabilize the three-dimensional structure. Mutations are performed on the hypervariable residues of the three CDR regions as listed in Table 1.

The mutation strategy is as follows: Alignment of a large number of nanobodies shows that certain residues in the CDR domains are variable, referred to as hypervariable residues. The hypervariable residue positions, and the occurrence probabilities of the 20 amino acids for each hypervariable position are identified by Kruse et al.^18^ In the present study, point mutations are applied on the twelve hypervariable positions, starting from Alanine up to Valine for each position, by using visual molecular dynamics (VMD).^19^ After each mutation, the nanobody is subjected to a minimization, annealing, conventional MD run and another minimization cycle by using NAMD.^20^ The interaction energy between the nanobody and the protein is then calculated, which contains electrostatic, VdW and non-bonded terms. The amino acid that leads to lowest energy mutation is selected. This is repeated independently for each mutation site. The resulting sequences are given in Table 1 along with the corresponding sequence of the wild type.

**Table 1.** Hypervariable CDR residues of EgA1 in 4KRO and their mutations. First row indicates the CDR region, second shows the residues of the wild type, and the next two rows show their mutations.

| CDR | 1 | 1 | 1 | 1 | 2 | 2 | 3 | 3 | 3 | 3 | 3 | 3 |
| --- | --- | --- | --- | --- | --- | --- | --- | --- | --- | --- | --- | --- |
| Residue | 30 | 31 | 32 | 33 | 53 | 54 | 99 | 100 | 101 | 102 | 103 | 104 |
| 4KRO | SER | SER | TYR | ALA | TRP | SER | THR | TYR | LEU | SER | SER | ASP |
| 4KROm1 | GLN | PHE | LYS | PRO | GLN | GLU | ASP | ARG | ASP | GLU | ARG | ARG |
| 4KROm2 | GLU | LYS | LYS | ALA | TRP | GLU | ASP | ARG | ASP | GLN | ARG | ARG |

### The Gaussian Network model

The GNM ^16^ is based on determining the number of spatial neighbors of a given residue that lie within a sphere of a given distance. This calculation is repeated for all of the N residues. The distance, referred to as the cutoff radius, *r*_*C*_, is generally taken between 7.0 and 7.2 Å.^16^ The latter is chosen here. The connectivity or the Kirchoff matrix, Γ, is obtained from the calculated distances as:

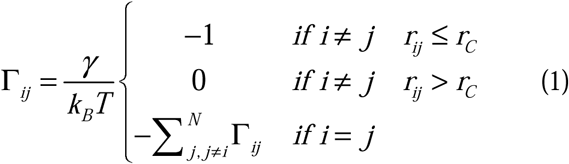

where *r*_*ij*_ is the distance between residues i and j, *γ* is a proportionality constant referred to as the spring constant of a virtual spring that represents the interaction between two neighboring residues, *k*_*B*_ is the Boltzmann constant and *T* is the absolute temperature. In all calculations here, we take 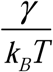 as unity because its actual value is immaterial for the purpose of this paper.

### Calculation of distances between residue pairs

As a common practice, the distances are calculated from the coordinates of atoms given in the Protein Data Bank crystal structure of proteins. In some cases, as in 4KRO for example, the data set lacks coordinate information on certain residues. For this reason, we use the following alternative strategy: We run molecular dynamics simulations of the protein in water and record all coordinates of the alpha carbons during a period where the protein has reached equilibration and find the distance between all alpha carbon pairs. Here we used 2000 frames where frames are separated by 0.01 ns. We then form the Γmatrix for each frame, and take its average. This leads to a Γmatrix with fractional entries and not 0’s and 1’s.

### Calculation of correlations and B-factors

According to the theory, the correlation between fluctuations of atom i and j is obtained from the inverse of the Kirchoff matrix as

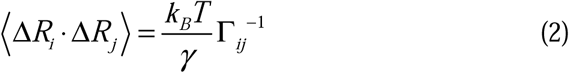

Here, Δ*R*_*i*_ is the vector that represents the instantaneous fluctuation of residue i and the dot represents the scalar product. The angular brackets denote the average of the dot products over time. By definition, the Kirchoff matrix is singular. A convenient way of forming the inverse is by calculating the singular values of the Kirchoff matrix. The formulation is made much clearer by using the eigenvectors of the Kirchoff matrix as basis vectors. In this representation, correlations are obtained in terms of the eigenvalues and eigenvectors as

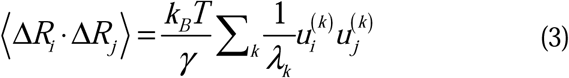

where, *λ*_*k*_ is the k^th^ eigenvalue and 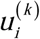 is the i^th^ component of k^th^ eigenvector. More detailed explanation of the eigenvector representation may be found in references ^16, 17, 21, 22^.

Equating j to i in Eq 3. gives the mean-square fluctuations of the i^th^ atom

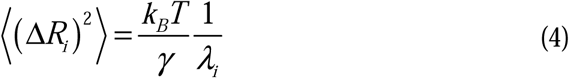

When a residue in a protein is replaced by another, the physicochemical properties of the new one may not be suitable for that location and may be exerting extra forces on other residues around it. Similarly, when a residue interacts with the surroundings, there may be forces on it. We denote correlations in the presence of perturbation of residue i by *A*_*ij*_(*τ*) = ⟨Δ*R* (0)· Δ*R*_*j*_(*τ*)⟩. In the Appendix, we show that a random perturbation on the i^th^ residue leads to time dependent correlations with the remaining residues according to:

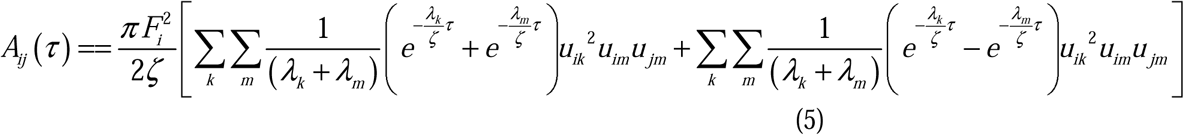

This equation reflects the dynamic aspect of perturbation-response: Firstly, it vanishes when the force vanishes. It is the correlation set in excess of correlations of the unperturbed structure under white noise (see Reference ^22^ for the derivations under white noise). The first double sum represents the synchronous and the second represents the asynchronous contribution to perturbation-response. The two terms are analyzed separately below. If *τ* is large, residue j will forget the effects of the perturbation. The matrix is not symmetric, i.e., *A*_*ij*_(*τ*) ≠ *A*_*ji*_(*τ*), which shows that the effect of perturbing residue i on j is not the same as the effect of perturbing j on i. The duration of perturbation on a responding residue may be defined by a correlation time. The following expression gives the correlation time 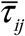 which is a measure of the time of decay of perturbation of residue j when it is perturbed by residue i at *τ* = 0 :

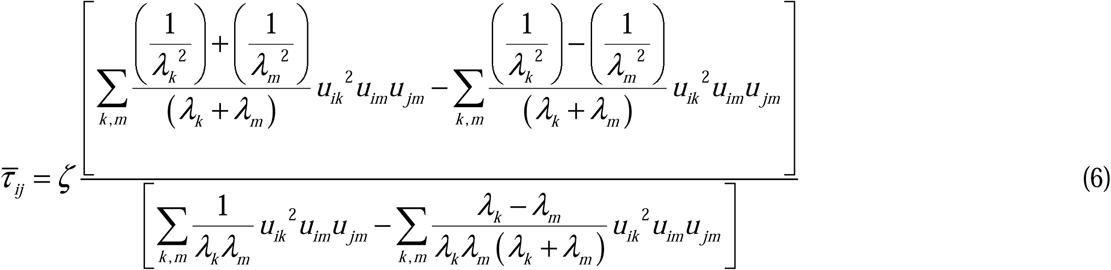

The first term in the numerator of Eq. 6 is the synchronous contribution to response time. The second is the asynchronous contribution. Similar to the perturbation-response correlations, the correlation time matrix is not symmetric, the effect of perturbing i on j may be of a different duration than the effect of perturbing j on i.

## Results

### Changes in B-factors upon mutation

The B-factors for the wild type 4KRO calculated by Eq 4. are presented on the left panel of Figure 2

**Figure 2.**
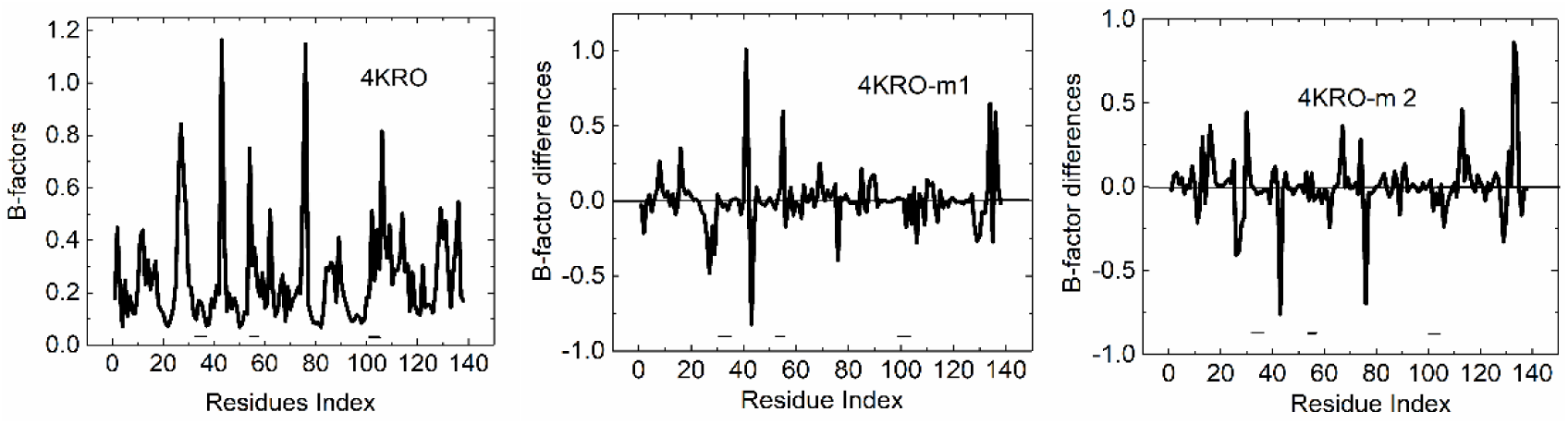
B-factors for the Nanobody/VHH domain EgA1 in 4KRO.pdb (left panel). The ordinate values of the middle panel are obtained as the difference between the B-factors of m1 and those of the wild type. Similarly, B-factor differences for m2 are shown in the right panel. Ordinate values are in Å^2^. CDR residues are indicated by the three horizontal bars at the lower part of each panel.

The front factor in Eq. 1 is taken as unity, therefore the abscissa values do not correspond to the actual B-factor values. The three horizontal dashed lines show the locations of the three CDRs where mutations are performed. In the left panel, the dashed lines show that residues of the first CDR do not fluctuate much, the second and third CDRs exhibit larger fluctuations. Largest fluctuations are observed in regions outside the three CDRs: ARG27, LYS43, LYS76, and SER106 are the most significant ones. The B-factor differences between the mutated proteins and the wild type are shown in the second and third panels. Each of the latter two panels is obtained by subtracting the B-factors of the wild type protein from those of the mutated one. When compared with the B-factors of the left panel the differences in B-factors resulting from mutation are significant and in the order of B-factors. It is interesting to note that mutations cause changes in parts of the nanobody that are outside the three CDR regions. LYS43 and LYS76 peaks are common in both m1 and m2, showing that these residues are most affected by mutation. Apart from changes common to m1 and m2, there are changes such as fluctuations of PRO41 which show significant increase for m1 but not affected much in m2. Similarly, ARG113 is affected in m2 but not in m1.

### Changes in correlations upon mutation

The correlations are shown in the three panels of Figure, the left panel is for the wild type and the center and right ones for the modified nanobodies. Comparison of the three panels clearly shows the effect of mutation on residue pair correlations. Certain patterns in the wild type are lost upon mutation. The wild type nanobody, left panel, shows that ARG27, LYS43, LYS76, and SER106 are correlated with several other residues. These systematic correlations are lost, however, upon mutation in both m1 and m2. New correlations are established in the middle and right panels of Figure 3, mostly in the tail of the protein. If we regard the presence of systematic correlations as a requirement for function, then loss of these correlations by mutation becomes important. Among correlations shown for the wild type, the ones which LYS76 is involved are worth discussing. LYS76 makes three hydrogen bonds with THR28 in the wild type, which are lost upon mutation. The absence of the LYS76-THR28 interaction changes the stability of the system of beta strands and all correlations of LYS76 are lost in a zipper-like fashion as may be seen from Figure 3. Similar to LYS76, correlations of ARG27, LYS43, and SER106 are also lost in the mutated structures.

**Figure 3.**
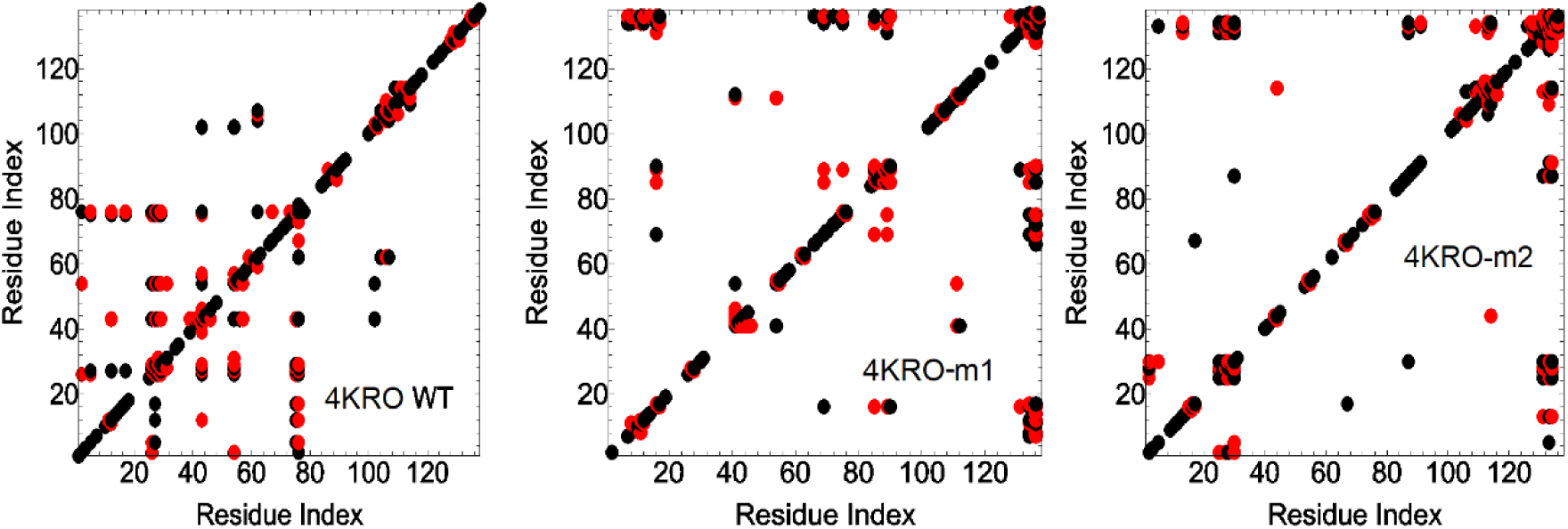
Residue correlations in wild type, m1 and m2. Red circles show negative and black circles show positive correlations.

### Effects of perturbing a residue and changes upon mutation

Response of nanobodies to perturbation is presented in Figure 4. Values of the ratio *A*_*ij*_(*τ*) / ⟨Δ*R*_*i*_·Δ*R*_*j*_⟩ is plotted in each panel. The numerator values are obtained from Eq. 5 and the corresponding denominator values are from Eq. 3. The abscissae show the residues that are perturbed and the ordinates show residues that respond. In Eq. 5 with 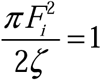 and 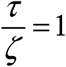. Values change between 0 and 0.32. For clarity, 20% of the points with largest ratio values are shown. The vertical pattern of points for the wild type in the left panel shows that certain residues (ARG27, LYS43, LYS76, and SER106) affect large numbers of other residues. This set of four residues are the ones that exhibit the largest correlations with other residues in the unperturbed system, as may be seen by comparison with Figure 3. The patterns observed for the wild type are diminished for the modified nanobodies. Interestingly, mutations create new responding residues at the tail of the nanobody.

**Figure 4.**
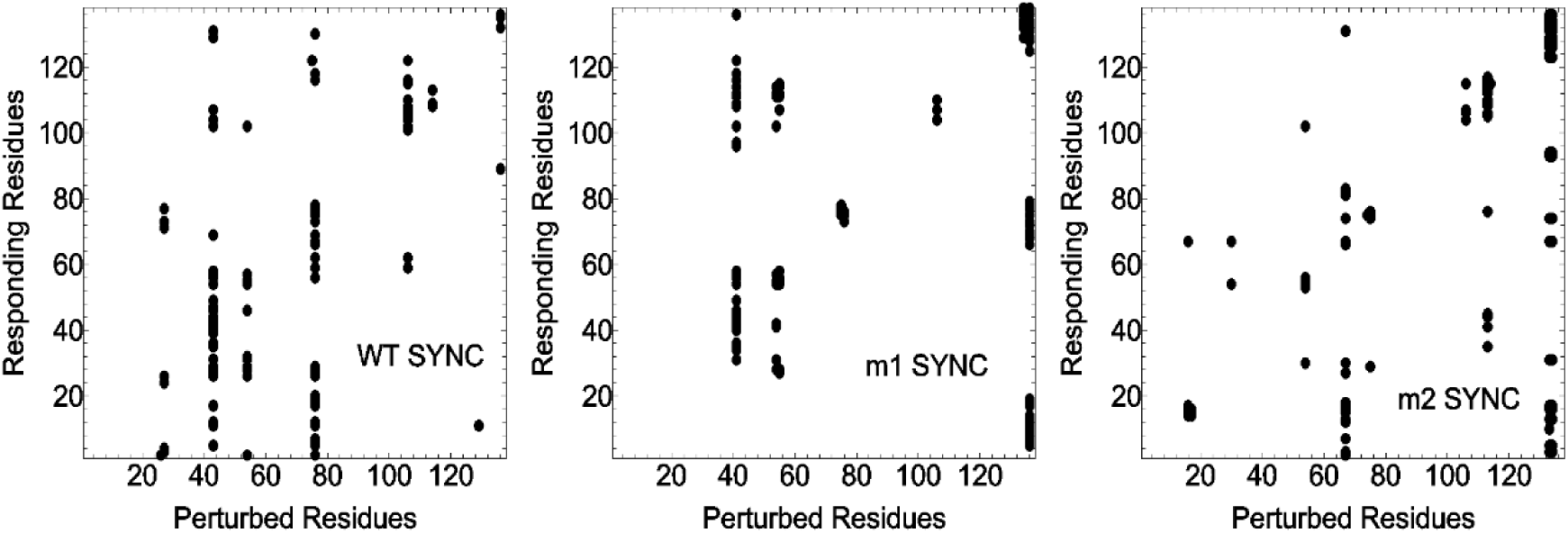
Synchronous residue responses in wild type (left panel), and modified nanobodies (central and right panels) to random perturbations. Only 100 points with largest responses (top 20%) are shown.

Ratio of asynchronous response to correlations is always negative, showing that asynchronicity always decreases response to perturbation. The asynchronous perturbation-response plots are presented in Figure 5 for the wild type (left panel), and the modified ones (center and right panels). For the wild type and m1, residue LYS43 is dominant, whose perturbation diminishes the response of several residues. For m2, this pattern is lost. Response values vary between 0 and -0.1. For clarity, only the top 20% of points are shown on the plots.

**Figure 5.**
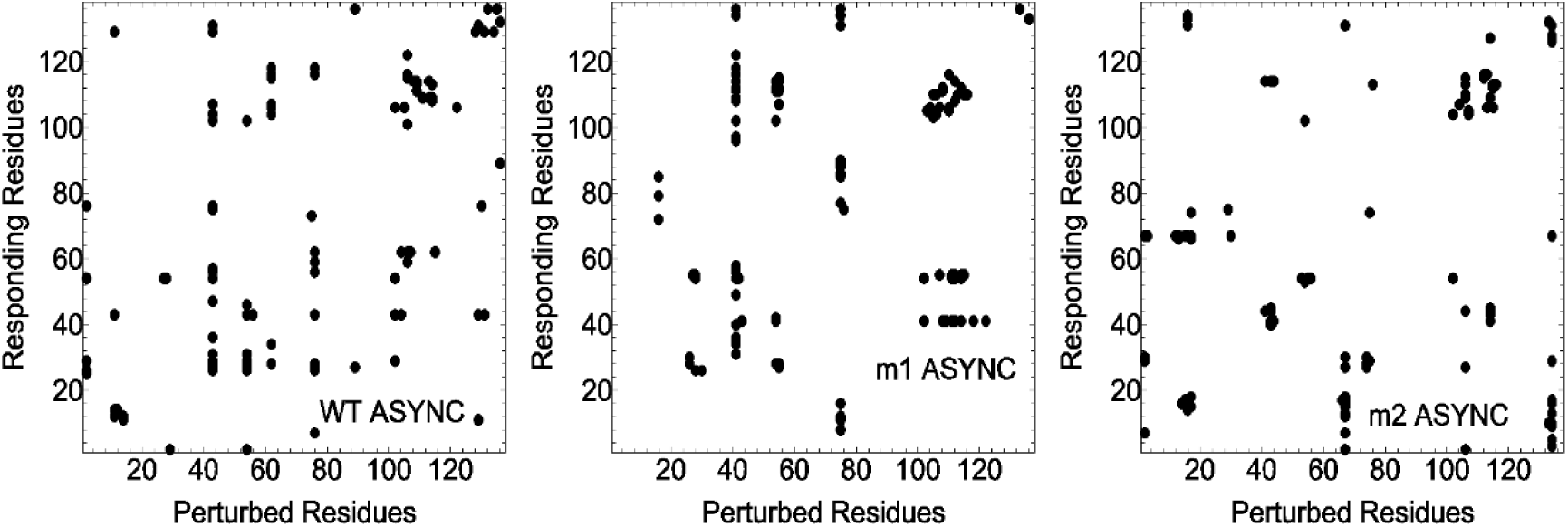
Asynchronous residue responses in wild type (left panel), and modified nanobodies (central and right panels) to random perturbations. Only 100 points with largest responses (top 20%) are shown.

When a residue is perturbed, its effect on another residue persists for some time and the duration of this effect may be important. Figure 6 shows how long this effect survives for the three samples studied. The left figure for the wild type shows three horizontal bands, for three residues, ARG27, LYS43, and LYS76. These three residues are the responding ones for which perturbation of several residues show their effect on these for long times. The correlation times vary between 0 and 30. Only the top 20% of the points are shown in the figures. Compared with results for the wild type, the three well defined horizontal lines have shifted to other locations in the modified nanobodies. Tail residues in the modified samples feel the effect of perturbation most strongly.

**Figure 6.**
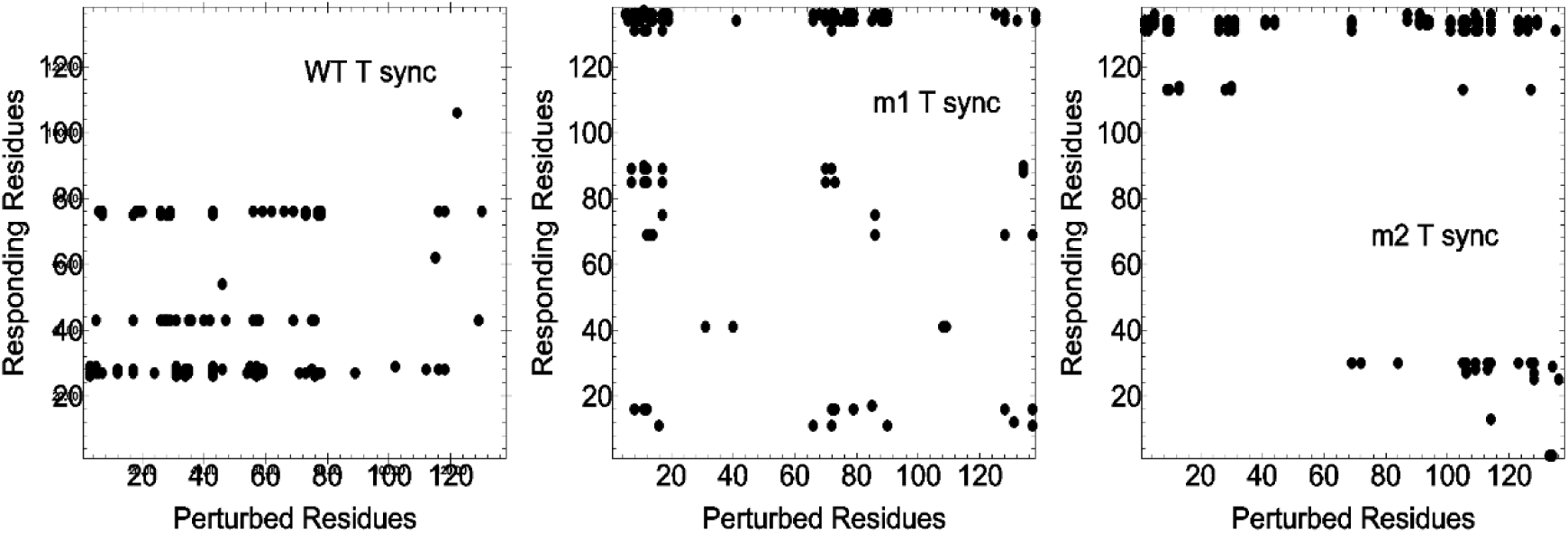
Residues with long synchronous response time in wild type (left panel), and modified nanobodies (central and right panels) to random perturbations. The correlation times vary between 0 and 30. Points representing the top 20% of correlation times are shown.

## Discussion and Conclusions

Using a simple harmonic model, we show that conformational features and dynamics of nanobodies change significantly upon mutation of the CDR residues. Correlation and perturbation-response patterns and in response times exhibit changes in information transfer features in the protein upon mutations. For each mutation site, the residue giving the lowest intermolecular energy of the system was chosen. Our calculations show that certain correlation patterns that exist in the nanobody produced in nature are lost upon mutation. However, as may be seen from results in the Supplementary Material, new patterns may set upon mutation. Whether these patterns are the determinants of good binders or not is not known. We are only reporting the changes observed.

The perturbation-response plots give interactions between several pairs of residues. These pairs are mostly spatial neighbors but there are spatially distant pairs as well. The pattern of perturbation-synchronous response shown in Figure 4 is similar to the correlation pattern of Figure 3. Perturbation of residues ARG27, LYS43, LYS76, and SER102 results in the strongest response of several other residues of the nanobody. Perturbing ARG27 leads to a strong response of LYS43 which is at a distance of 33 Å. LYS43 operates on VAL12, SER54 and SER102 which are 31 and 33 and 31 Å apart, respectively. SER54 affects GLN1 which is at a distance of 21 Å. All of these interactions are missing in the mutated nanobodies m1 and m2. Mutation leads to new interactions, however. The dominant ones emerging after mutation are made with the participation of tail residues. (See also the changes shown in Supplementary Material.

Values of synchronous and asynchronous components of *A*_*ij*_(*τ*) for 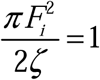 and 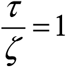 for each residue responding to the perturbation of ARG27 are presented in Figure 8. The curves show that asynchronous response always opposes synchronous response and diminishes the synchronicity in the system. Mutation decreases the responses as may be seen by comparing the ordinate values of the central and right panels with that of the wild type, left panel. Synchronicity is a dynamic property that shows the correlations among residue pairs as a function of time. It is a simple measure of information transfer in proteins. A more realistic measure is entropy transfer which is based on conditional residue pair probabilities rather than simple pair correlations. Nonzero pair correlations between two residues simply indicates that information may be transferred between the two. Similar to entropy transfer, *A*_*ij*_(*τ*) is directional also, indicating the degree of causality in the perturbed and responding system. Since asynchronicity always decreases synchronicity, it should be regarded as an unwanted property of the system as far as information transfer is concerned. Furthermore, it is nonhomogeneous, affecting some residue pairs different than others.

**Figure 7.**
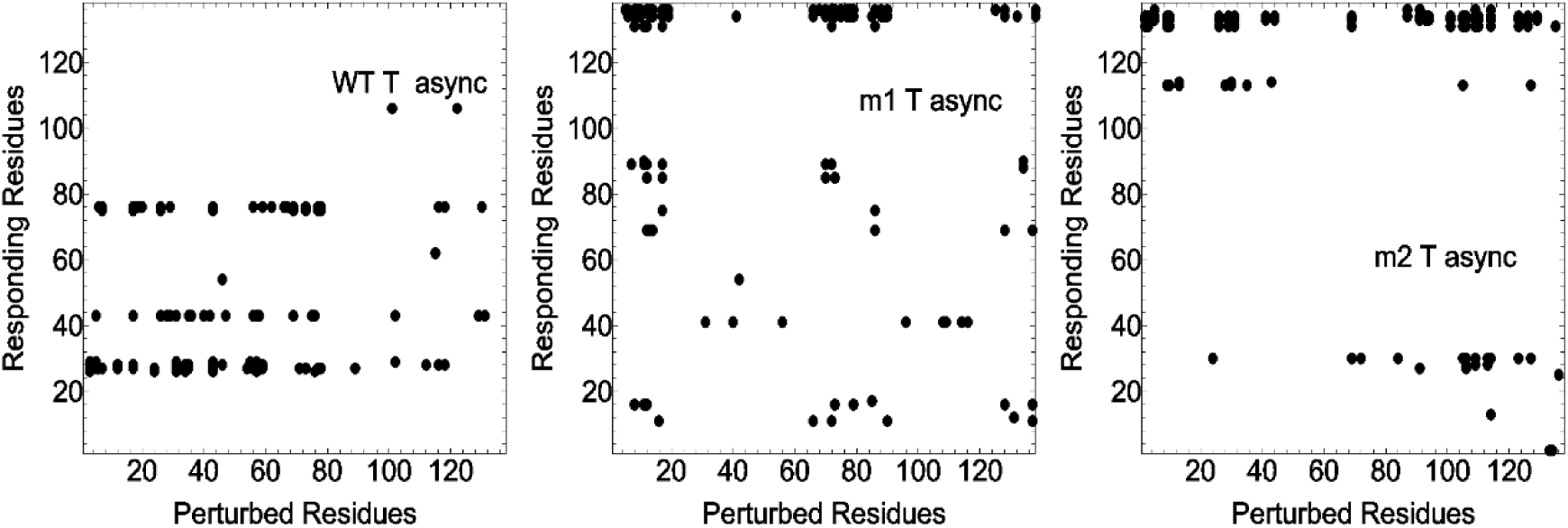
Residues with long asynchronous response time in wild type (left panel), and modified nanobodies (central and right panels) to random perturbations. The correlation times vary between 0 and 30. Points representing the top 20% of correlation times are shown.

**Figure 8.**
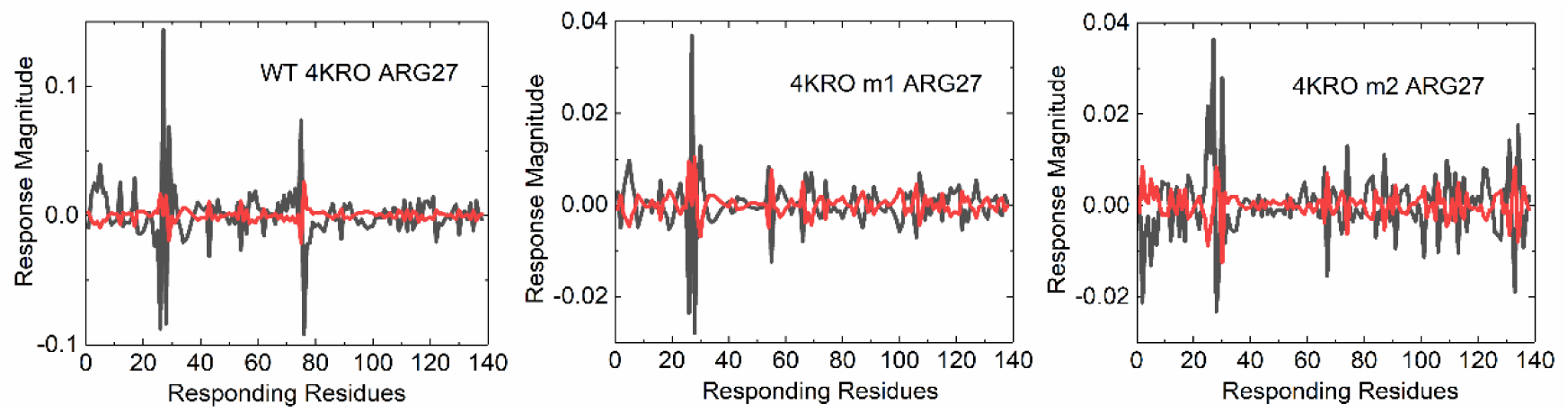
Synchronous and asynchronous response magnitudes of residues to perturbation of ARG27. Black line is for synchronous, red line is for asynchronous responses. Left panel is for wild type, central and right panels are for mutated samples m1 and m2.

While nanobodies are used as examples in the present study, the formulation developed is general and is applicable to any protein where the dynamics of perturbation-response is of interest. Effects resulting from the mutation of a residue may be accepted as mimicking allostery effects where the force field around the mutated element is perturbed and response of other residues are investigated. The theory developed focuses on the change of fluctuations in a protein rather than change of mean coordinates. Role of fluctuations in protein behavior, especially in allostery, is important. Recently, there has been a paradigm change in the understanding of allostery, where allostery without conformational change emphasizing the role of fluctuations both static and dynamic came under focus.^23, 24^

## Appendix

### Synchronicity and asynchronicity

The behavior of a harmonic system in viscous medium and in the presence of noise obeys the Langevin equation:

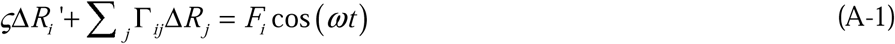

where, the prime is the time derivative, *ζ* is the friction coefficient, and *F*_*i*_ cos(*ωt*) is the external force acting on the i^th^ atom. The solution of this equation leads to the fluctuation of residues, Δ*R*(*t*), as

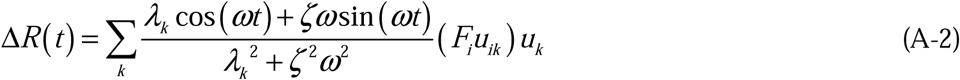

Here, Δ*R*(*t*) is a column vector whose entries show the displacement of each atom. The derivation for the expression for Δ*R*(*t*) is given in Reference ^22^. The time delayed correlation of residue fluctuations is defined as:

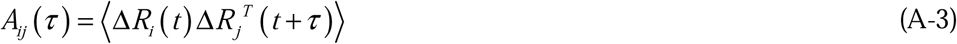

Here, the angular brackets denote average over the full trajectory of motion, the superposed T represents the transpose of the vector and *A*_*ij*_(*τ*) is the time delayed correlation of fluctuations between residues i and j where the i^th^ atom is excited at time t by a perturbing effect and the residue j responds at time *t* +*τ*.

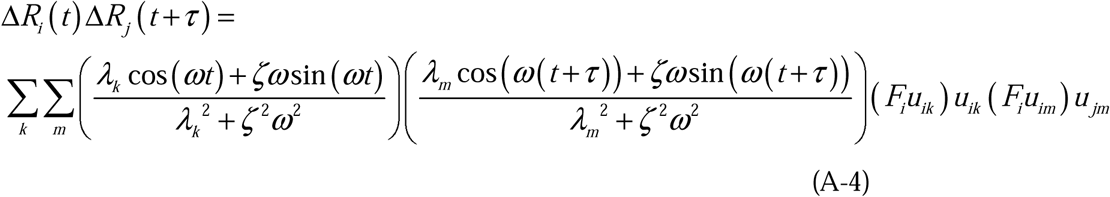

The term *u*_*ik*_ is the i^th^ element of the kth eigenvector, with similar definitions for the others. When residue i is perturbed by a cosine function, its correlation with another residue i a time *τ* after the excitation, averaged over time, t, is

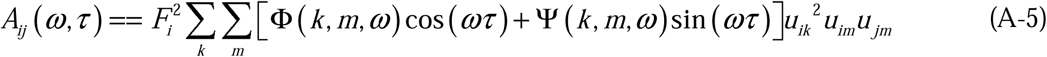

where, Φ(*ω*) and Ψ(*ω*) are the synchronous and asynchronous components, respectively:

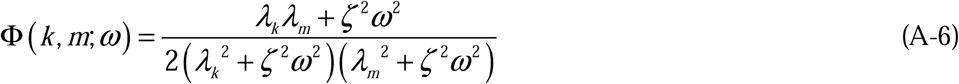

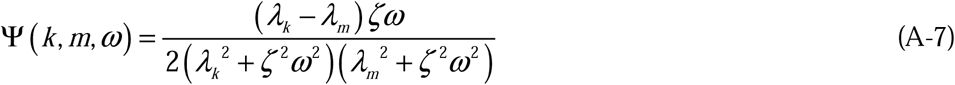

In random excitation, the cosine perturbation will not be at a single frequency, but will have a continuum of frequencies. We therefore integrate *A*_*ij*_(*ω,τ*) over all possible frequencies as follows:

Using the residue theorem, the integrals over frequencies become

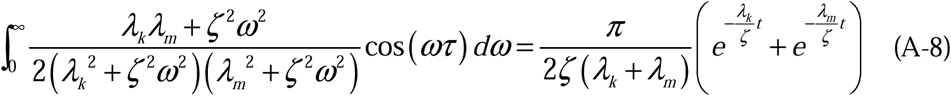

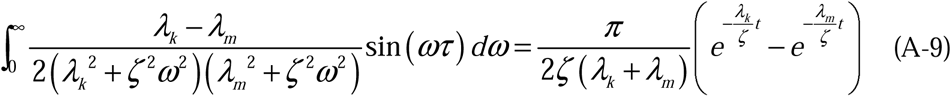

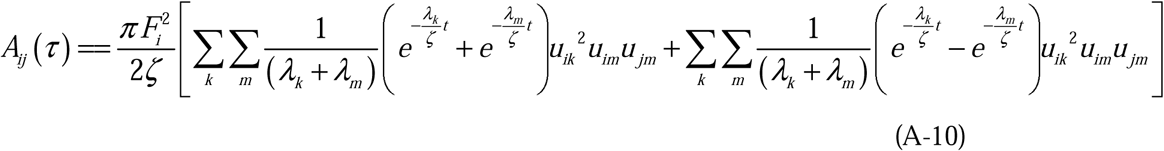

The correlations *A*_*ij*_(*τ*) vanish when there is no force. Therefore, these correlations are deviations from the unperturbed correlations of the protein. When t=0, the asynchronous component equates to zero. Similarly, the response of an atom to its own perturbation does not contain an asynchronous term because the term *u*_*ik*_^2^*u*_*im*_ *u*_*jm*_ = *u*_*ik*_^2^*u*_*im*_^2^ becomes symmetric with respect to k and m.

It will be of interest to know how long the effects of a perturbation last. A correlation time for the decay of a perturbation effect may be defined as

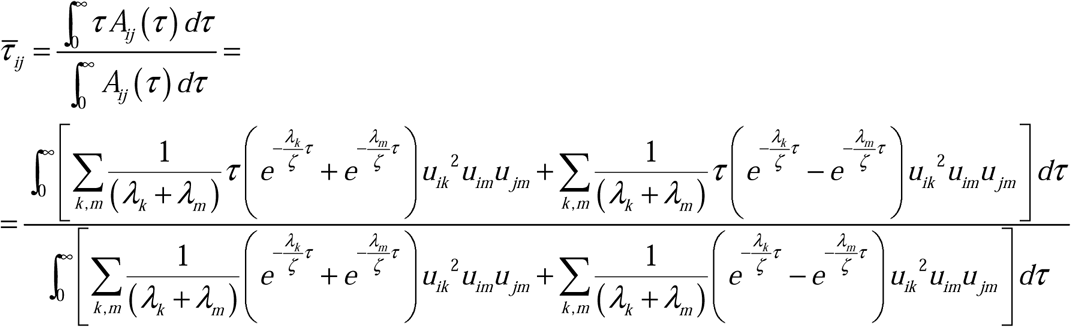

Performing the integrations, we obtain

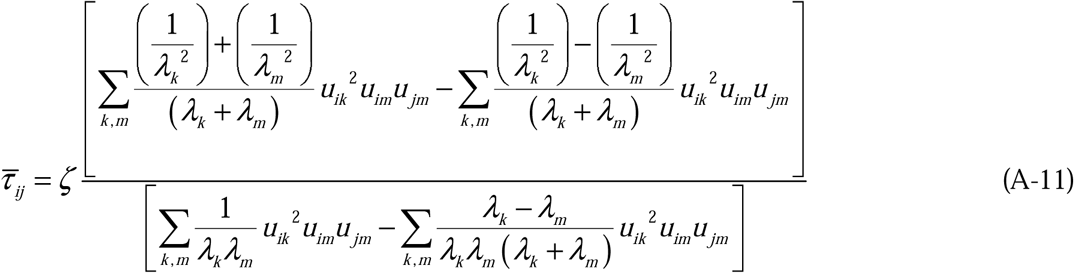

