## Supplemental Material for "Fluctuation, correlation and perturbation-response behavior of nature-made and artificial nanobodies"

## 4I0C

Table S-1. Hypervariable CDR residues of 4I0C and their mutations. First row indicates the CDR region, second shows the residues of the wild type, and the next row shows their mutations.

| CDR | 1 | 1 | 1 | 1 | 1 | 2 | 2 | 3 | 3 | 3 | 3 | 3 | 3 | 3 | 3 |
|---|---|---|---|---|---|---|---|---|---|---|---|---|---|---|---|
| Residue | 27 | 28 | 29 | 30 | 31 | 55 | 56 | 97 | 98 | 101 | 110 | 111 | 112 | 113 | 115 |
| 4I0C | LEU | SER | THR | THR | VAL | PHE | PRO | LYS | THR | PHE | SER | ARG | ALA | TYR | HIS |
| 4I0Cm1 | ASN | GLN | VAL | GLU | TRP | ASN | ASP | ALA | ILE | GLY | TRP | SER | LYS | TRP | LYS |

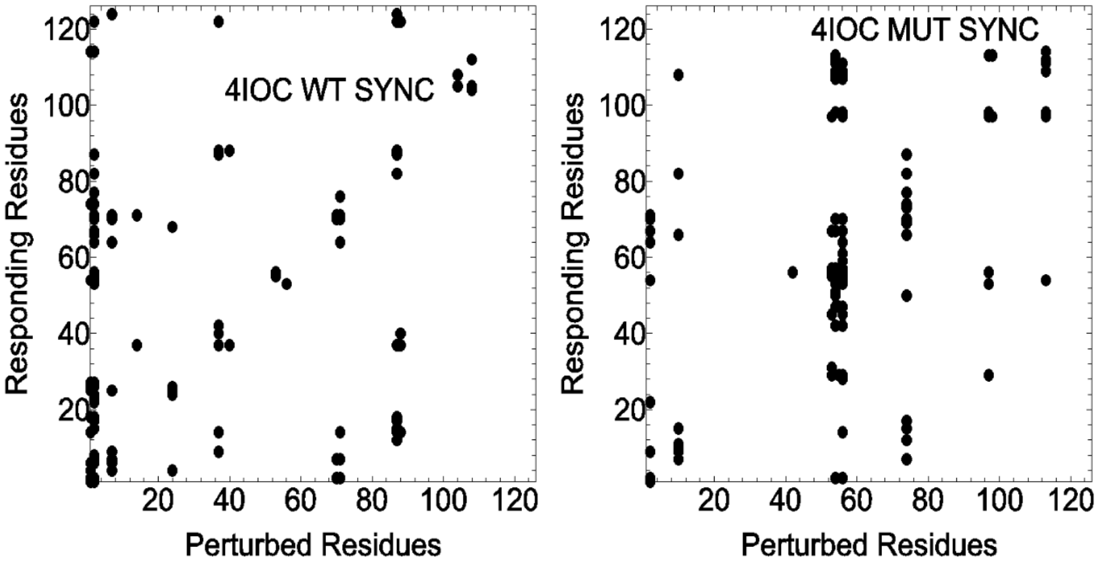


Figure S-1 Synchronous residue responses to perturbation in wild type 4I0C (left panel) and mutated 4I0C (right panel). Points represent top 15% of responses.

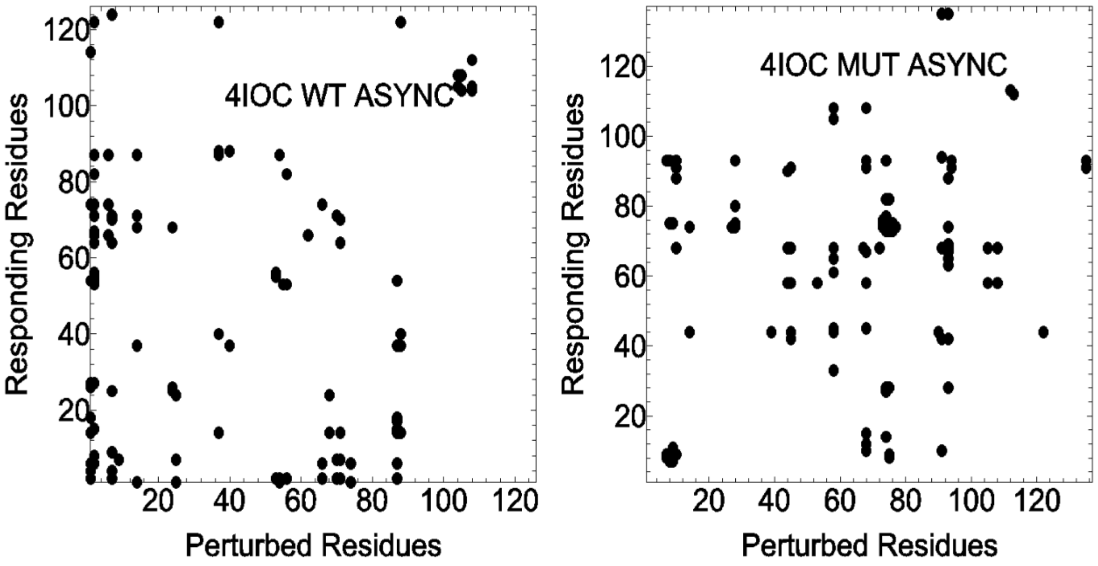


Figure S-2 Asynchronous residue responses to perturbation in wild type 4I0C (left panel) and mutated 4I0C (right panel). Points represent top 15% of responses.

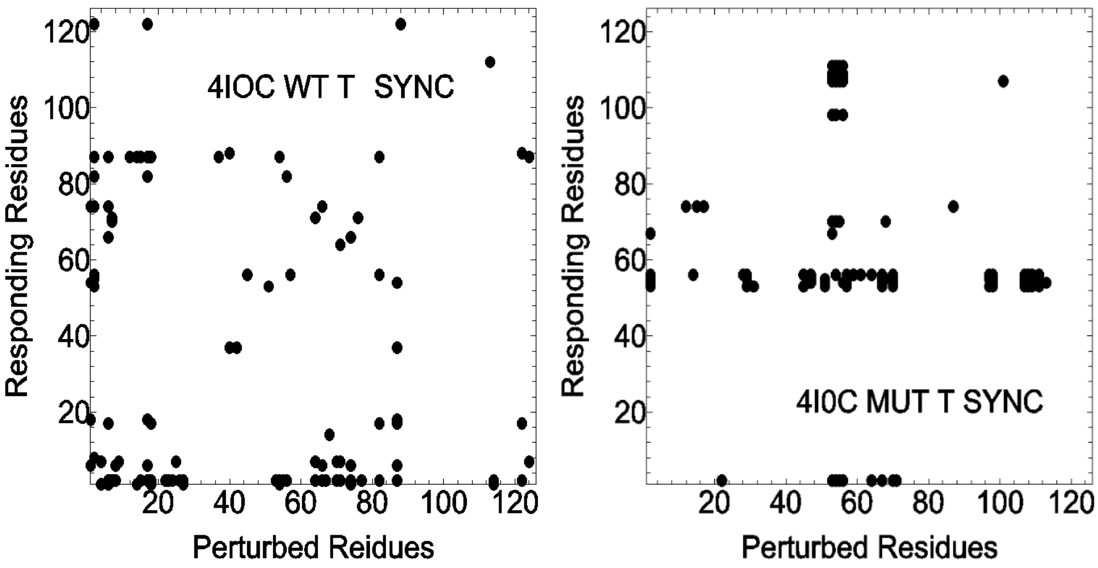


Figure S-3 Residues with long synchronous response time to perturbation in wild type 4I0C (left panel) and mutated 4I0C (right panel). Points represent top 15% of responses.

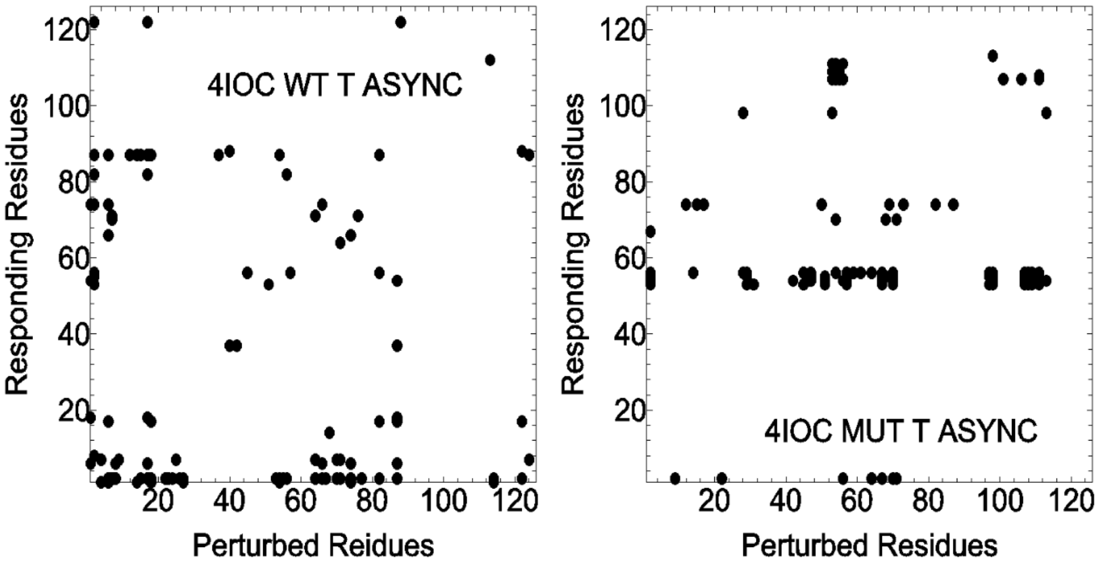


Figure S-4 Residues with long asynchronous response time to perturbation in wild type 4I0C (left panel) and mutated 4I0C (right panel). Points represent top 15% of responses.

# 5O0W

Table S-2. Hypervariable CDR residues of 5O0W and their mutations. First row indicates the CDR region, second shows the residues of the wild type, and the next row shows their mutations.

| CDR | 1 | 1 | 1 | 2 | 3 | 3 | 3 | 3 | 3 | 3 | 3 | 3 | 3 |
|---|---|---|---|---|---|---|---|---|---|---|---|---|---|
| Residue | 28 | 31 | 32 | 53 | 103 | 104 | 105 | 106 | 108 | 110 | 115 | 125 | 126 |
| 5O0W | ALA | TYR | TYR | ARG | ASP | THR | THR | ASP | TYR | SER | TYR | ASP | TYR |
| 5O0Wm1 | HIS | SER | GLN | LEU | ALA | ILE | ALA | THR | THR | GLY | ALA | GLU | ARG |

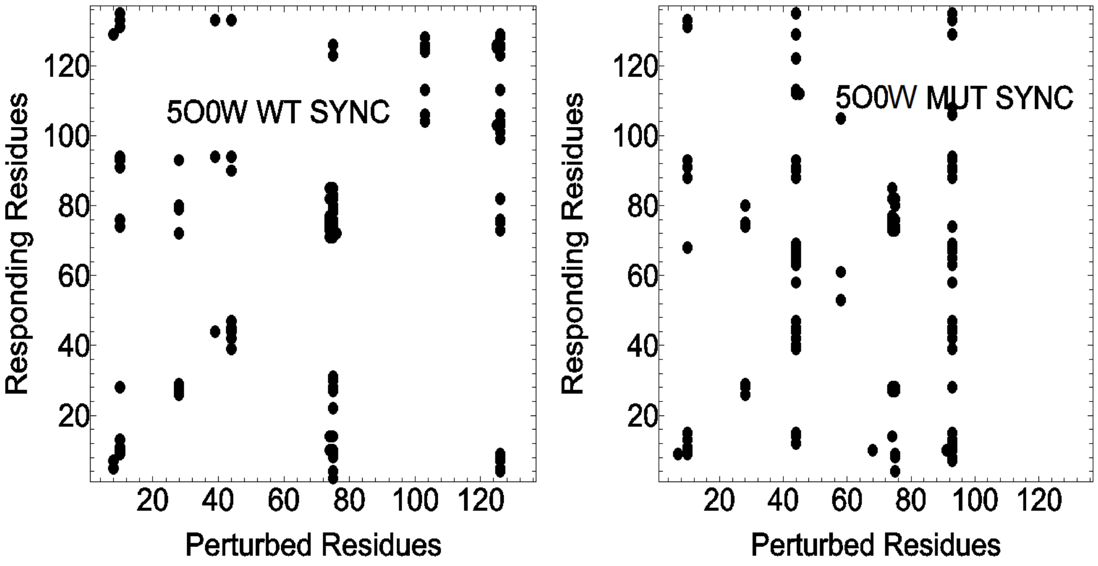


Figure S-5 Synchronous residue response to perturbation in wild type 5O0W (left panel) and mutated 5O0W (right panel). Points represent top 15% of responses.

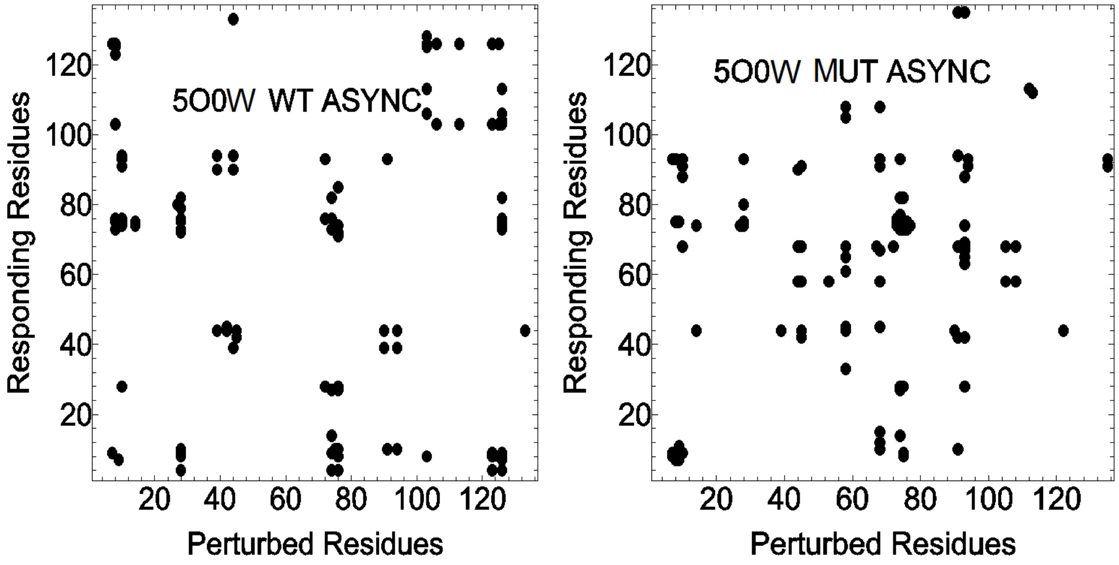

Figure S-6 Asynchronous residue response to perturbation in wild type 5O0W (left panel) and mutated 5O0W (right panel). Points represent top %15 of responses.

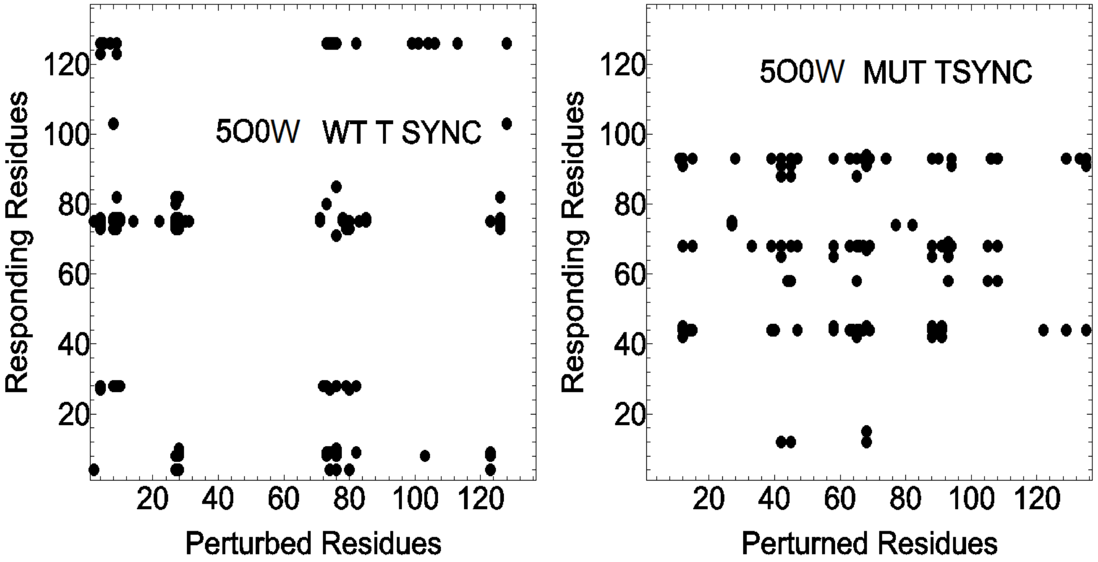


Figure S-7 Residues with long synchronous response time to perturbation in wild type 5O0W (left panel) and mutated 5O0W (right panel). Points represent top 15% of responses.

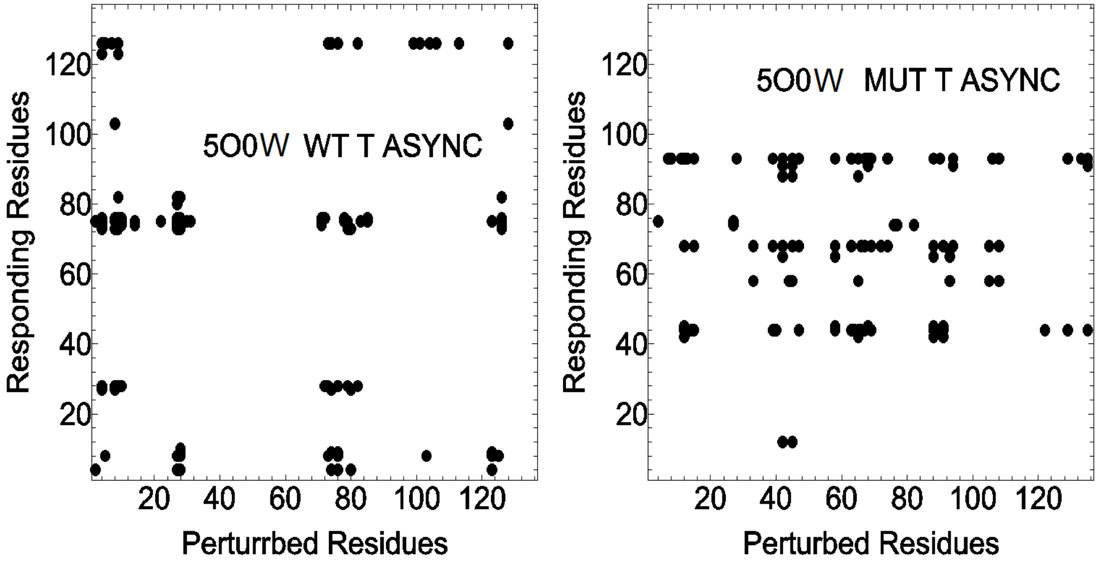


Figure S-8 Residues with long asynchronous response time to perturbation in wild type 5O0W (left panel) and mutated 5O0W (right panel). Points represent top 15% of responses.
